# Radiosensitivity is an acquired vulnerability of PARPi-resistant BRCA1-deficient tumors

**DOI:** 10.1101/370577

**Authors:** Marco Barazas, Alessia Gasparini, Yike Huang, Asli Küçükosmanoğlu, Stefano Annunziato, Peter Bouwman, Wendy Sol, Ariena Kersbergen, Natalie Proost, Marieke van de Ven, Jos Jonkers, Gerben Borst, Sven Rottenberg

**Author notes:** Corresponding authors (J.J), (G.B.), (S.R.).

## Abstract

The homologous recombination (HR) defect in BRCA1-associated cancers can be therapeutically exploited by the treatment with DNA-damaging agents and poly (ADP-ribose) polymerase (PARP) inhibitors. We and others previously reported that BRCA1-deficient tumors are initially hypersensitive to the inhibition of topoisomerase I/II and PARP, but acquire drug resistance through restoration of HR activity by the loss of end-resection antagonists of the 53BP1/RIF1/REV7/Shieldin/CST pathway. Here, we identified radiotherapy as an acquired vulnerability of 53BP1;BRCA1-deficient cells *in vitro* and *in vivo*. In contrast to the radioresistance caused by HR restoration through BRCA1 reconstitution, HR restoration by 53BP1 pathway inactivation further increases radiosensitivity. This highlights the relevance of this pathway for the repair of radiotherapy-induced damage. Moreover, our data show that BRCA1-mutated tumors that acquired drug resistance due to BRCA1-independent HR restoration can be targeted by radiotherapy.

**STATEMENT OF SIGNIFICANCE:** In this study, we uncovered radiosensitivity as a novel therapeutically exploitable vulnerability of BRCA1-deficient mouse mammary cells that have acquired drug resistance due to the loss of the 53BP1 pathway.

## INTRODUCTION

Most of the currently used anti-cancer therapies include applications that target the DNA such as topoisomerase inhibitors, DNA-crosslinking agents and radiotherapy. In recent years, it has become clear that alterations in the DNA damage response (DDR) provide a useful explanation for the initial drug sensitivity. Most cancers have lost a critical DDR pathway during cancer evolution (1,2), and therefore respond to clinical interventions that cause DNA damage. To further exploit defects in the DDR, targeted therapies have been developed using the “synthetic lethal” approach (3). Tumors that have lost specific DDR pathways rely more heavily on the remaining pathways, while normal tissues still have all DDR pathways available. Thus, inhibition of a critical backup pathway in DDR-deficient cells will cause lethality in tumor cells while not harming the normal cells. A prime example is the selective toxicity of poly(ADP-ribose) polymerase inhibitors (PARPi) to cancer cells that are defective in homologous recombination (HR) due to dysfunctional BRCA1/2 proteins (4). Indeed, PARPi provide an opportunity to achieve a major benefit for patients with HR-deficient cancers, if the hurdle of drug resistance can be overcome (4). Besides resistance mechanisms that involve restoration of BRCA1/2 protein function, there are a number of BRCA1-independent roads to PARPi resistance. Most notably, we and others have found that the loss of end-resection antagonists of the 53BP1/RIF1/REV7/SHLD/CST DNA repair pathway partially restores HR activity and causes PARPi resistance in BRCA1-deficient cells (5-10). In this study, we demonstrate that these PARPi-resistant tumor cells show increased radiosensitivity. This finding was spurred by our initial observation that, in contrast to PARPi-resistance, acquired radioresistance in *K14cre;Brca1^F/F^;p53^F/F^* (KB1P) mouse mammary tumors with irreversible deletions in *Brca1* was not mediated by the loss of 53BP1, nor by restoration of HR. Further *in vitro* and *in vivo* examination of the genetic interaction between BRCA1 and the 53BP1 pathway on therapy response established radiosensitivity as an acquired vulnerability of KB1P tumor cells that have inactivated the 53BP1 pathway and thereby provides insight in new treatment strategies to target PARPi-resistant tumors.

## MATERIALS & METHODS

### In vivo studies

All animal experiments were approved by the Animal Ethics Committee of The Netherlands Cancer Institute (Amsterdam, the Netherlands) and performed in accordance with the Dutch Act on Animal Experimentation (November 2014).

Radiosensitivity responses were evaluated by allografting previously harvested tumor pieces derived from the *K14cre;Trp53^F/F^* (KP) and *K14cre;Brca1^F/F^;Trp53^F/F^* (KB1P) genetically engineered mouse model (11). The tumor volume was determined using the egg formula (length x width^2^ x 0.5). Established tumors (>500 mm^3^) were irradiated daily using a high-precision small-animal irradiator equipped with a cone-beam CT scanner (X-RAD 225Cx). The dosing schedule consisted of 36Gy/9f in 3 weeks. Radioresistant tumors were generated by allografting KB1P tumor pieces in 6-9 week old syngeneic female mice followed by daily treatment with 2, 4 or 8Gy, until a predetermined response was achieved at which point the treatment was halted. The treatment was reinitiated when the tumor relapsed to the starting volume, and this was repeated until the tumor eventually stopped responding (KB1P-RR). KB1P-RR tumors were harvested and collected in formalin or DMSO for downstream analysis.

The stability of radioresistance and cross-resistance profiles were determined by allografting KB1P-RR and matched treatment-naïve (KB1P-N) tumor pieces in 6-9 week old syngeneic female mice. Radiotherapy was given to established tumors (>500 mm^3^) and consisted of 36Gy/9f in 3 weeks. The cross-resistance study was carried out on established tumors (>200 mm^3^), at which point mice were stratified into the different treatment arms. Treatments consisted of olaparib (50 mg/kg drug i.p. on 28 consecutive days (12)), topotecan (4 mg/kg drug i.p. on days 0-4 and 14-18 (13)), cisplatin (6 mg/kg drug i.v. single dose (12)) or untreated. To assess the radiotherapy response in isogenic *Trp53bp1*-knockout or control tumors (sg*NT*), previously generated and validated tumor organoids were allografted in 6-9 week old NMRI nude female mice, as described (14). Briefly, tumor organoids were collected, incubated with TripLE at 37°C for 5’, dissociated into single cells, washed in PBS, resuspended in tumor organoid medium and mixed in a 1:1 ratio of tumor organoid suspension and BME in a cell concentration of 10^4^ cells per 40 μl. Subsequently, 10^4^ cells were transplanted in the fourth right mammary fat pad of 6-9 week-old NMRI nude female mice. Treatment of tumor bearing mice was initiated when tumors reached a size of 50-100 mm^3^, at which point mice were stratified into the untreated or radiotherapy treatment group. The dosing schedule consisted of 40Gy/10f in 2 weeks. Animals were sacrificed with CO_2_ when the tumor reached a volume of 1,500mm^3^.

### Tumor analysis

Immunohistochemistry to detect 53BP1 expression in tumor tissues was performed as described (7). RAD51 immunofluorescence on tumors was performed as described (8). Briefly, ionizing-radiation induced foci (IRIF) were induced by irradiation (15 Gy) 2 hours prior to tumor harvesting and fixation in formalin. The staining was performed on FFPE slides using a non-commercial antibody for RAD51 provided by R. Kanaar (1:5,000) and Goat anti-Rabbit IgG (H+L) Cross-Adsorbed Secondary Antibody, Alexa Fluor 568 from Thermo Fisher Scientific, catalog # A-11011, RRID AB_143157 and mounted and counterstained using using Vectashield mounting medium with DAPI (H1500, Vector Laboratories). Samples were imaged by confocal microscope (Leica SP5, Leica Microsystems GmbH) using a 63x oil objective. Multiple different Z-stacks were imaged per sample. The number of foci per nucleus was analyzed automatically using an ImageJ script (8).

### Cell culture

The KB1P-G3 tumor cell line was previously established from a *K14cre;Brca1^F/F^;Trp53^F/F^* (KB1P) mouse mammary tumor and cultured as described (7). Briefly, cells were cultured in DMEM/F-12 medium (Life Technologies) in the presence of 10% FCS, penicillin/streptomycin (Gibco), 5μg/mL insulin (Sigma), 5ng/mL epidermal growth factor (Life Technologies) and 5ng/mL cholera toxin (Gentaur) under low oxygen conditions (3% O_2_, 5% CO^2^ at 37°C). The KB1PM7-N 3D tumor organoid lines were previously established from a *Brca1^-/-^;p53^-/-^;Mdr1a/b^-/-^* mouse mammary tumor and cultured as described (14).Briefly, KB1PM7-N 3D tumor organoid cells were seeded in Basement Membrane Extract Type 2 (BME, Trevigen) on 24-well suspension plates (Greiner Bio-One) and cultured in AdDMEM/F12 supplemented with 1 M HEPES (Sigma), GlutaMAX (Invitrogen), penicillin/streptomycin (Gibco), B27 (Gibco), 125 μM N-acetyl-L-cysteine (Sigma), 50 ng/mL murine epidermal growth factor (Invitrogen).

### Plasmids and genome editing

Unless otherwise stated, CRISPR/SpCas9-targeted KB1P-G3 tumor cell lines were generated using a modified version of the pX330 backbone (15) in which a puromycin resistance ORF was cloned under the hPGK promoter (16). sgRNA sequences were cloned in the pX330puro backbone as described (15) and sequence verified by Sanger sequencing. KB1P-G3 tumor cells were transfected using TransIT-LT1 (Mirus) reagents according to manufacturer’s protocol and using conditions as described (10). Briefly, 150,000 cells were seeded in 6-well plates one day prior to transfection with 1 μg DNA. 24 hours later, the medium was refreshed with puromycin (3 μg/mL) containing medium and cells were selected for three days. KB1P-G3 BRCA1 reconstituted cells were generated by transfecting KB1P-G3 cells with a human *BRCA1* cDNA expression construct (17) using Lipofectamine 2000 (Thermo Fisher Scientific). One day after transfection, cells were passaged and cultured with 300 μg/ml G418 (Thermo Fisher Scientific) to select for human *BRCA1* complemented colonies. G418 resistant colonies were tested for human *BRCA1* integration by PCR with human *BRCA1* exon 11 specific primers: Fwd-5’-TCCAGGAAATGCAGAAGAGG-3’, Rv-5’- ACTGGAGCCCACTTCATTAG-3’.

Lentivirus production in HEK293T cells was performed as described (18). KB1P-G3 cells were transduced with the FUCCI plasmids mKO2-hCdt1(30/120) and mAG-hGem(1/110) (19) and were sorted for red fluorescence and green fluorescence in two subsequent sorting rounds to ensure the presence of both constructs in each cell. Cell lines targeted with the pGSC_Cas9_Neo and pLenti-sgRNA-tetR-T2A-Puro system (20) were generated by lentiviral transduction. sgRNA sequences were cloned in the pLenti-sgRNA-tetR-T2A-Puro backbone using the BfuAI (NEB) restriction enzyme. Following transduction and selection with puromycin (3 μg/mL) and blasticidin (500 μg/mL) for five days, cells were cultured in the presence of 3 μg/mL doxycycline (Sigma) for at least five days to induce sgRNA expression. Doxycycline was removed from the medium prior to starting the competition assay. The KB1PM7-N 3D tumor organoid line derivatives KB1PM7-N sg*NT* and KB1PM7-N sg*Trp53bp1* were established previously (14).

sgRNA sequences used in this study are (14): sg*Trp53bp1*: 5’- GAACAATCTGCTGTAGAACA-3’, sg*Trp53bp1-2*: 5’-TACCGGGCTGTACTGTAACA-3’, sg*Rif1*: 5’-GACAATCCTGAGGTAATTCT-3’, sg*Rev7*: 5’- GCGCAAGAAGTACAACGTGC-3’ and sg*Ctc1*: 5’_CTTGAAGCCGAACAGTGCCA-3’. Targeting efficiencies were determined by genomic DNA isolation (Gentra Puregene, Qiagen) followed by PCR amplification of the target loci and subsequent Sanger sequencing. Sequences were analyzed using the TIDE algorithm (21), using parental cells as a reference. The following PCR primers were used: sg*Trp53bp1*: Fwd-5’- TGAGAAATGGAGGCAACACCA-3’ and Rv-5’-CTCGATCTCACACTTCCGCC-3’, sg*Trp53bp1-2*: Fwd-5’-GAGAGCGCACGCACAGTAAG-3’ and Rv-5’- TGGGCTGGCTCTGATACTTTG-3’, sg*Rif1*: Fwd-5’- GACGGACGCCTACCTAACTC-3’ and Rv-5’-AAAGGCCCTTGACATCTAGCC-3’, sg*Rev7*: Fwd-5’- TAGCCCGGTCGTAGATTGGA-3’ and Rv-5’-CTGTCCGCTATCAGCCTCTG-3’, and sg*Ctc1*: Fwd-5’-TGTTCCAGACAGGGATTTTCCAA-3’ and Rv-5’- AGGAGAGGGTTGCTTCAGGA-3’.

### Western blotting

Western blotting for 53BP1, RIF1 and REV7 was performed as described (8). Briefly, cells were plated in equal amounts and the next day cells were washed with PBS and lysed with RIPA lysis buffer supplemented with protease inhibitors. The protein concentration was determined using the Pierce BCA Protein Assay Kit (Thermo) according to manufacturer’s protocol. Samples were heated at 70°C for 10 minutes and loaded on 3-8% Tris-acetate (53BP1 and RIF1) or 4-12% Bis-Tris gradient gels (REV7). Following SDS-PAGE separation, proteins were transferred to nitrocellulose membranes for 53BP1 and RIF1 (Invitrogen), or to PVDF membrane (Millipore) for REV7. Membranes were blocked in 5% milk with Tris-buffered saline Triton X-100 buffer (100mMTris, pH7.4, 500mMNaCl, 0.1% TritonX-100) (TBS-T0.1%) and all antibody incubations were performed in the same buffer. Primary antibody incubations were performed overnight at 4°C, secondary antibody incubations were performed for 1 hour at RT and proteins were visualized by ECL. Antibodies used: rabbit anti-53BP1 (ab21083, Abcam), 1:1,000 dilution; rabbit anti-RIF1 (SK1316, (6)); mouse anti-REV7 (612266, BD Biosciences), 1:5,000 dilution; mouse anti-a-tubulin (T6074, Sigma), 1:5,000 dilution; polyclonal rabbit anti-mouse immunoglobulins/HRP (P0161, Dako), 1:10,000 dilution; polyclonal swine anti-rabbit immunoglobulins/HRP (P0217, Dako), 1:10,000 dilution.

### *In vitro* drug-response profiles

Clonogenic growth assays with PARPi (olaparib and AZD2461), topoisomerase inhibitors (topotecan and doxorubicin) and the alkylating agent cisplatin were performed as described (10). Briefly, on day 0, 5 x 10^3^ KB1P-G3 cells were seeded per 6-well and drugs were added in the indicated concentrations. Untreated cells were fixed on day 6 and treated cells were fixed on day 9. After fixation, cells were stained with 0.1% crystal violet and plates were scanned with the Gelcount (Oxford Optronix). Crystal violet was solubilized using 10% acetic acid and the absorbance at 562nm was measured using a Tecan plate reader. The experiment was performed three times. Dose-response curves with PARPi were generated similarly, but 1 x 10^3^ KB1P-G3 cells were seeded per 6-well and all conditions were fixed on day 9. Radiotherapy survival curves were generated by seeding different amounts of cells (50, 100, 500, 1000 and 5000) per 6-well in technical duplicates, followed by irradiation with a single fraction of the indicated dose using the ^137^Cs-irradiation unit Gammacell 40 EXACTOR (Best Theratronics). Plates were fixed and stained with 0.1% crystal violet on day 9. Colonies containing at least 50 cells were counted manually using an inverted microscope, selecting the wells in which <150 colonies were counted to restrict the quantifications to wells in which the colonies were still well-separated. Plating efficiencies (PE) were calculated by dividing the number of colonies after treatment with the amount of cells that were originally plated. Surviving fractions were calculated for each KB1P-G3 cell line by dividing the PE after RT to the PE of untreated cells and plotted using Graphpad Prism. Data were fitted to LQ model: Y=exp(-(A^*^x + B^*^x^2)). The experiment was performed at least two times.

Competition assays were performed as described (10). Briefly, KB1P-G3 SpCas9 expressing cells were transduced with pLenti-sgRNA-tetR-T2A-Puro vectors in which the indicated sgRNAs were cloned. Following selection with puromycin (3 μg/mL) for three days and recovery from selection, 5,000 cells were plated in 6-well plates in triplicate per condition, with or without olaparib (75 nM), AZD2461 (250 nM) or radiotherapy (3fr/4Gy/1wk). After 10 days of treatment, cells were harvested, counted and re-plated at 5,000 cells per 6-well two times (total treatment time of 30 days). Cells were harvested for gDNA isolation and target loci analysis at day 0 and after treatment.

Growth curves were generated as described (10). KB1P-G3 cells were seeded at 1,000 cells per well in 96-well plates. In each experiment, 6 technical replicates were seeded and the well confluency was recorded every 4 hours using an IncuCyte Zoom Live – Cell Analysis System (Essen Bioscience). The images were analyzed using IncuCyte Zoom software. The experiment was performed three times.

### Quantification and statistical analysis

Statistical analysis was performed using Graphpad Prism software and R software. Significance was calculated as follows: unpaired two-tailed students t-test (Fig. 1B, 1C, 4B and Supplementary Fig. S1A), CFAssay Bioconductor version 3.7 (Fig. 2A,C), Kruskal Wallis non-parametric test (Supplementary Fig. 2D), Log-Rank (Mantel-Cox) test (Fig. 4D and Supplementary Fig. S1B), ordinary one-way ANOVA with Dunnett’s multiple comparison test (Supplementary Fig. 4B), Binomial test (Supplementary Fig. 2B), using observed distributions of PARPi-resistant KB1P(M) tumors for comparison (17; data not shown).

**Figure 1.**
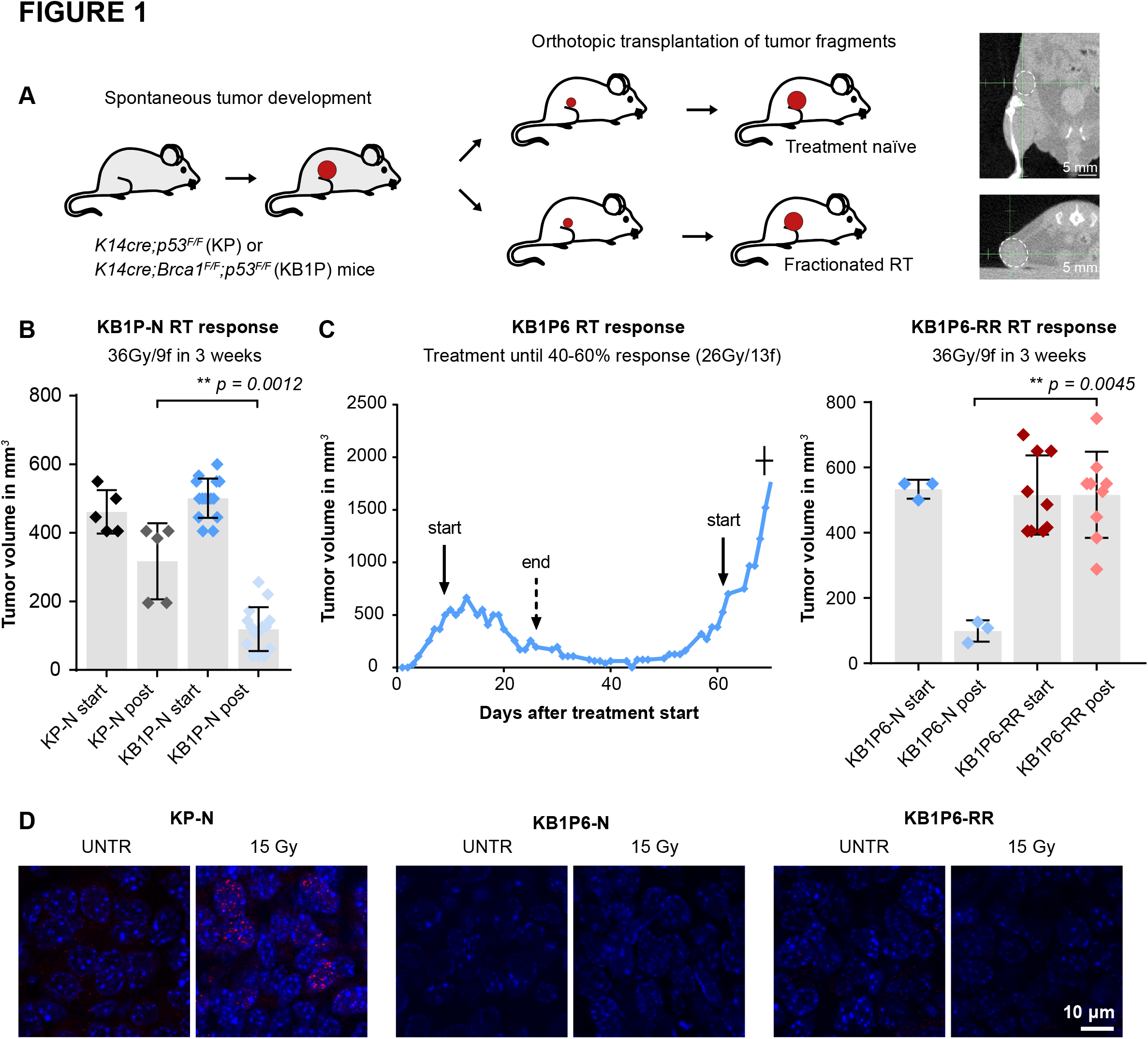
Radioresistance in KB1P tumors is not accompanied by HR restoration. See also Supplementary Figure 1 and Supplementary Figure 2. **A.** Schematic overview of the serial transplantation model to test the radiotherapy sensitivity. Tumors derived from the spontaneous KP or KB1P tumor model were allografted in syngeneic FVB mice and treated with fractionated radiotherapy when tumors reached 500 mm^3^. Radiotherapy was delivered locally using a high precision irradiator dedicated for mice and equipped with a cone-beam CT scanner. This allowed accurate localization and treatment of the tumor. Example images are shown with the tumor highlighted by a white dashed line. **B.** Radiotherapy response of KB1P compared to KP tumors. Radiotherapy was delivered to established tumors as 36Gy/9fr in 3 weeks. Tumor volumes were compared at the end of treatment. Data are plotted as mean ± SD. Significance was calculated by unpaired two-tailed students t-test. **C.** Example of a KB1P6 tumor in which radiotherapy resistance was induced by continuous treatment until 40-60% response was measured (13fr/2Gy). Treatment was reinitiated when the tumor had regrown to its starting volume and this was repeated until the tumor stopped responding (KB1P-RR). KB1P-RR and its matched KB1P-N tumors were allografted in syngeneic FVB mice and treated with fractionated radiotherapy when tumors reached 500 mm^3^ (36Gy/9fr in 3 weeks). The volume post treatment was compared to the volume at treatment start and are plotted as mean ± SD. Significance was calculated by unpaired two-tailed students t-test. **D.** Example immunofluorescence images of RAD51 IRIF formation on KP-N, KB1P-N and KB1P-RR tumors before or after 15Gy of IR.

**Figure 2.**
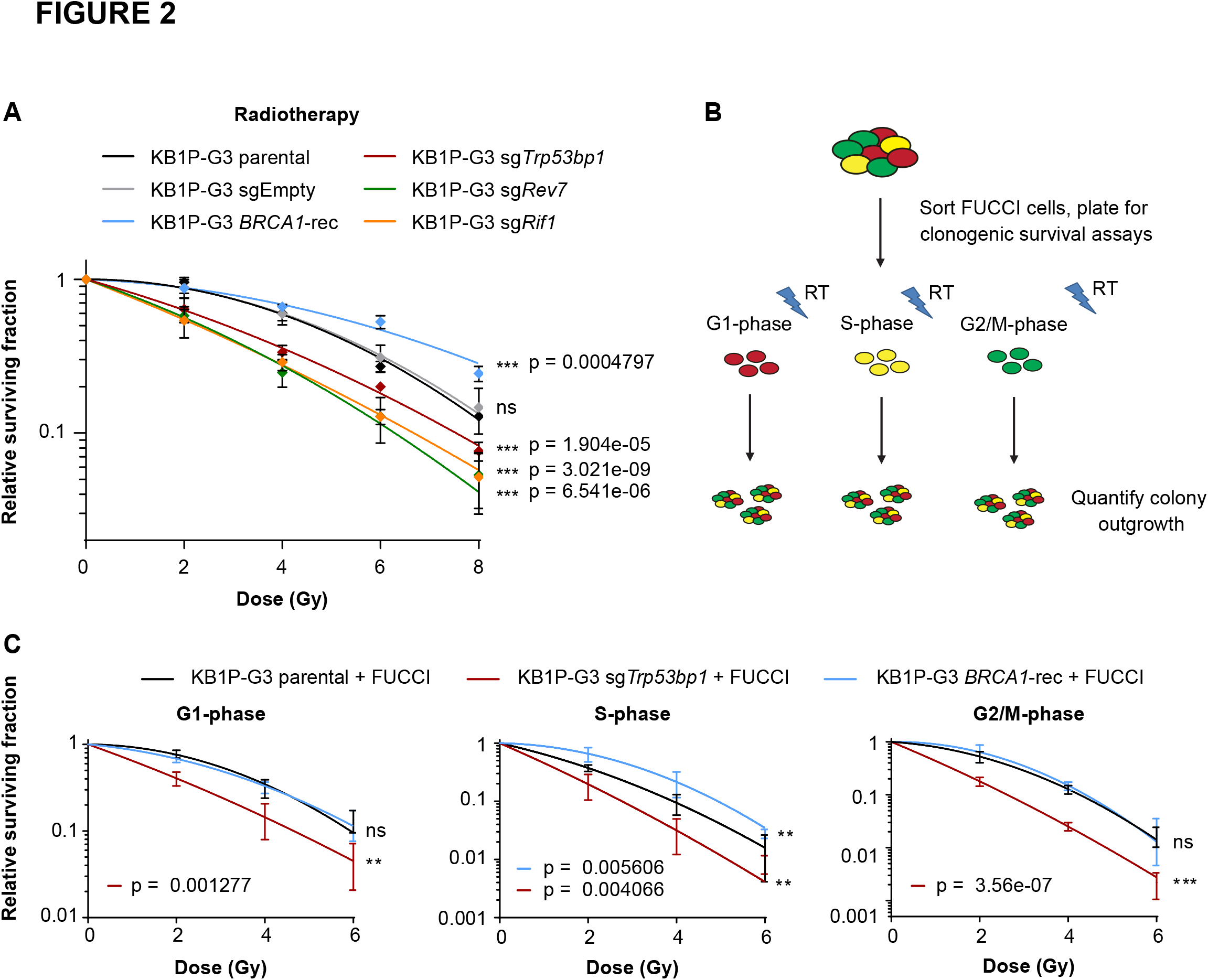
53BP1 pathway inactivation in KB1P-G3 tumor cells enhances radiosensitivity. See also Supplementary Figure 3 and Supplementary Figure 4. **A.** The radiotherapy response of CRISPR/spCas9 targeted KB1P-G3 tumor cell lines were determined by clonogenic survival assay. Cells were plated as single cells, irradiated with the indicated dose immediately after plating and fixed and stained 10 days later. The number of colonies was counted manually using an inverted microscope. Quantifications were performed blinded. Data represent at least two independent experiments and were plotted as mean ± SD and fitted to the LQ-model using GraphPad Prism software. Statistics were calculated using CFAssay in R. **B.** Schematic overview of the FUCCI experiment in which the radiotherapy response was analyzed per stage of the cell cycle. Cells were FACS sorted, plated and irradiated directly after plating. **C.** The radiotherapy response of sorted KB1P-G3 parental, sgTrp53bp1-targeted or BRCA1-reconstituted tumor cells in which the FUCCI system was introduced. Plates were fixed and quantified 10 days later as in Fig. 2A. Data represent three independent experiments and were plotted as in Fig. 2A. Statistics were calculated as in Fig. 2A.

## RESULTS

### KB1P TUMORS ACQUIRE RADIORESISTANCE INDEPENDENT OF HR RESTORATION

To test whether the loss of the 53BP1 pathway mediated radioresistance in the *K14cre;Brca1^F/F^;p53^F/F^* (KB1P) mouse model of hereditary breast cancer, we treated mice bearing KB1P mammary tumors with radiotherapy using a high-precision small-animal irradiator equipped with a cone-beam CT scanner (Fig. 1A) (22). This enabled us to deliver focused radiotherapy to the tumor whilst sparing normal tissue and to apply clinically relevant high dose radiotherapy regimens (e.g. 36Gy/9f in 3 weeks). Compared to treatment-naïve *K14cre;p53^F/F^* (KP-N) tumors, treatment-naïve KB1P-(KB1P-N) tumors were more sensitive to radiotherapy, highlighting the role of BRCA1-mediated DNA repair in response to radiotherapy (Fig. 1B). Resistance to radiotherapy was induced in KB1P-N tumors by halting treatment when either >90% or >40-60% tumor eradication was measured, followed by treatment re-initiation when tumors relapsed to the starting volume. This was repeated until tumors eventually stopped responding (Fig. 1C, Supplementary Fig. S1A). Notably, these radioresistant KB1P (KB1P-RR) tumors showed stable radioresistance, as serially transplanted KB1P-RR tumors were significantly less responsive to radiotherapy than their treatment-naïve counterparts (Fig. 1C, Supplementary Fig. S1A). The tumor takes- and growth rates of KB1P-RR and KB1P-N tumors were similar (data not shown). Interestingly, the KB1P-RR tumors were cross-resistant to olaparib and topotecan, which is indicative of enhanced DNA damage repair in KB1P-RR tumors (Supplementary Fig. S1B). Since these drug-response profiles are consistent with restored HR and resemble KB1P tumors that have lost 53BP1 (7,14), we analyzed 53BP1 expression by IHC in a panel of KB1P-RR tumors. Interestingly, while PARPi resistance was mediated by loss of 53BP1 expression in 12 out of 79 KB1P(M) tumors ((7); data not shown), all 26 KB1P-RR tumors retained 53BP1 expression (p = 0.0253, binomial test) (Supplementary Fig. S2A-B).

We next scored the capacity of KB1P-RR tumors to form ionizing radiation (IR)-induced foci of RAD51 as a measure of their HR status. Hereto, KB1P-RR tumors and their matched KB1P-N counterparts were orthotopically transplanted into syngeneic FVB/N mice. Established tumors (>1000mm^3^) were treated with 15 Gy RT and stained for RAD51 by immunofluorescence. We previously showed that HR restoration is frequently observed in PARPi-resistant KB1P tumors ((7,8); data not shown). Strikingly, none of 5 tested KB1P-RR tumors scored as RAD51-IRIF positive, demonstrating that restoration of HR was not a major resistance mechanism in response to radiotherapy (p = 0.0165, binomial test) (Fig. 1D, Supplementary Fig. S2C-D). Thus, BRCA1-deficient KB1P tumors, which have an increased radiosensitivity compared to BRCA1-proficient tumors, did not show HR restoration upon acquisition of radiotherapy resistance. This finding seems at odds with the observed partial HR restoration in PARPi-resistant tumors and is suggestive of an adverse effect of partial HR restoration on radiotherapy survival.

### BRCA1-INDEPENDENT RESTORATION OF HR INDUCES HYPERSENSITIVITY TO RADIOTHERAPY

To explore the radiotherapy response of BRCA1-deficient tumor cells harboring BRCA1-independent genetic events that restore HR, we focused on 53BP1. Drug response profiles were investigated in the KB1P-G3 tumor cell line in which *Trp53bp1* was knocked out using CRISPR/SpCas9 technology as evidenced by allele disruption and protein knockout (Supplementary Fig. S3A-B). KB1P-G3 tumor cells which were reconstituted with human BRCA1 (KB1P-G3 *BRCA1*-rec) were included as a control for HR proficiency. We measured the clonogenic growth of KB1P-G3-parental, KB1P-G3-sg*Trp53bp1*, KB1P-G3-*BRCA1*-rec and KB1P-G3-sgEmpty control cells in response to PARPi (olaparib, AZD2461), topotecan, doxorubicin and cisplatin. In line with previous studies (7,14), loss of 53BP1 or re-expression of BRCA1 induced resistance to these treatments (Supplementary Fig. S3C-D and S4). However, upon radiotherapy treatment (RT), depletion of 53BP1 resulted in a significant reduction in the surviving fraction compared to KB1P-G3-sgEmpty control cells. Similarly, depletion of *Rif1* or *Rev7/Mad2l2*, two downstream factors of 53BP1, also reduced the survival of KB1P-G3 cells upon RT treatment (Fig. 2A and Supplementary Fig. S3A-B). Although consistent with the absence of restoration of HR in KB1P-RR tumors, it was surprising to find that restoration of HR via loss of the 53BP1 pathway did not provide a survival benefit to RT. Indeed, in line with our findings that BRCA1-deficient tumors were more sensitive to radiotherapy compared to BRCA1-proficient tumors, restoration of HR by restoring BRCA1 expression in KB1P-G3 tumor cells did increase resistance to radiotherapy (Fig. 2A), underscoring the role of the HR repair pathway in response to RT treatment. Since repair via HR is restricted to DNA regions that undergo replication, we tested whether loss of 53BP1 might have opposing effects on cells in G1 versus cells in S/G2. Hereto, the FUCCI system was introduced in KB1P-G3-parental, KB1P-G3-sgEmpty, KB1P-G3-sg*Trp53bp1* and KB1P-G3-*BRCA1*-rec cells. Cells were plated and irradiated directly after fluorescence-activated cell sorting (FACS) to study clonogenic survival in response to RT as a function of each cell cycle phase (Fig. 2B). While reconstitution of BRCA1 enhanced the viability upon RT in S-phase populations, loss of 53BP1 sensitized KB1P-G3 tumor cells to RT in all phases of the cell cycle (Fig. 2C). These data show that the 53BP1 pathway plays an important role in the repair of RT-induced DNA damage throughout the cell cycle and that its loss in BRCA1-deficient cells is not compensated by the restoration of the HR pathway. Importantly, these data uncover a vulnerability of BRCA1-deficient cells that have restored HR via loss of the 53BP1-pathway.

To test whether radiotherapy can be applied to deplete PARPi-resistant cells within a mixed population of PARPi-resistant and -sensitive cells, we monitored the evolution of polyclonal populations of KB1P-G3 tumor cells targeted for *Trp53bp1*, *Rev7/Mad2l2*, *Rif1* and *Ctc1* (Fig 3A). The CST complex member CTC1 was included as we recently showed that loss of CTC1 restored HR and promoted PARPi resistance in BRCA1-deficient cells (10). KB1P-G3 tumor cells were transduced with an inducible CRISPR/Cas9 system to achieve approximately 50% modified alleles as quantified by Sanger sequencing followed by TIDE analysis (21). These polyclonal populations were subsequently plated for clonogenic growth and treated with olaparib, fractionated radiotherapy (3fr per week/4Gy/1wk) or left untreated. All populations were harvested after 10 days and re-plated three times in equal cell amounts, followed by quantification of the allele distribution of the surviving populations. As expected, PARPi-resistant populations were readily selected out upon targeting of members of the 53BP1 pathway or CTC1. The surviving populations mainly comprised cells enriched for disrupted alleles compared to untreated populations (Fig. 3B). In stark contrast, the abundance of targeted alleles in cells subjected to fractionated radiotherapy was markedly reduced compared to untreated populations, further underscoring the notion that restoration of HR through loss of the 53BP1 pathway creates a therapeutically exploitable vulnerability to radiotherapy.

**Figure 3.**
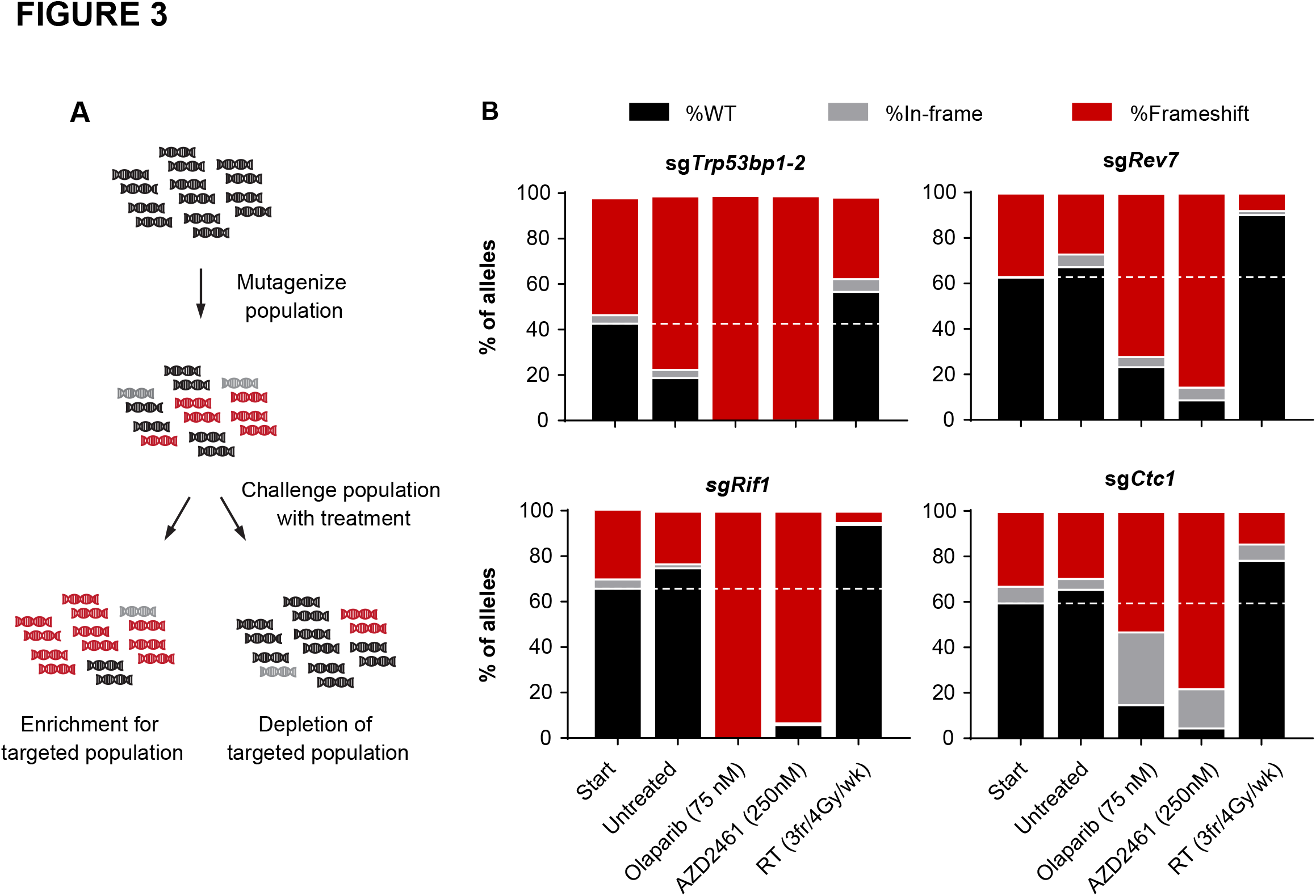
PARPi-resistant KB1P-G3 tumor cells can be depleted by radiotherapy. **A.** Schematic overview of the competition assay using polyclonal CRISPR/SpCas9-targeted starting populations. The allele distribution was quantified by TIDE before and after treatment with olaparib (75 nM), AZD2461 (250 nM) or IR (3fr/4Gy/wk). Cells were passaged and re-plated in equal cell amounts every 10 days for a total of 30 days. Untreated populations were taken along to assess the evolution in a neutral setting. **B.** TIDE quantifications of the indicated targeted populations at treatment initiation and after the last indicated treatment (day 30).

### LOSS OF 53BP1 IN KB1P MOUSE MAMMARY TUMOR-DERIVED ORGANOIDS ENHANCES THE SENSITIVITY TO RADIOTHERAPY *IN VIVO*

We next examined whether the therapeutic vulnerability exposed by 53BP1 loss is exploitable *in vivo* using our recently established tumor organoid model (14). Hereto, KB1PM7-N tumor-derived organoids were transduced with a control or a *Trp53bp1*- targeting sgRNA and subsequently allografted in mice. Fractionated radiotherapy consisting of 5 consecutive fractions of 4Gy per week for 2 weeks was initiated on mice bearing established tumors (50-100 mm^3^) and the effect on tumor volume was evaluated at the end of treatment (Fig. 4A). Hereby, the growth of KB1PM7-N sg*NT* tumors could be contained (Fig. 4B-C). While radiotherapy extended the survival of all treated mice, depletion of 53BP1 significantly enhanced the response, resulting in markedly reduced tumor volume at the end of treatment (p = 0.0081, t-test.) and a prolonged time to relapse, defined as five times the original treatment volume (Fig. 4D) (p = 0.0117, Log-rank test). Together, these data demonstrate that KB1P tumors that have lost 53BP1 expression are sensitized to radiotherapy, despite being unresponsive to PARPi treatment (14).

**Figure 4.**
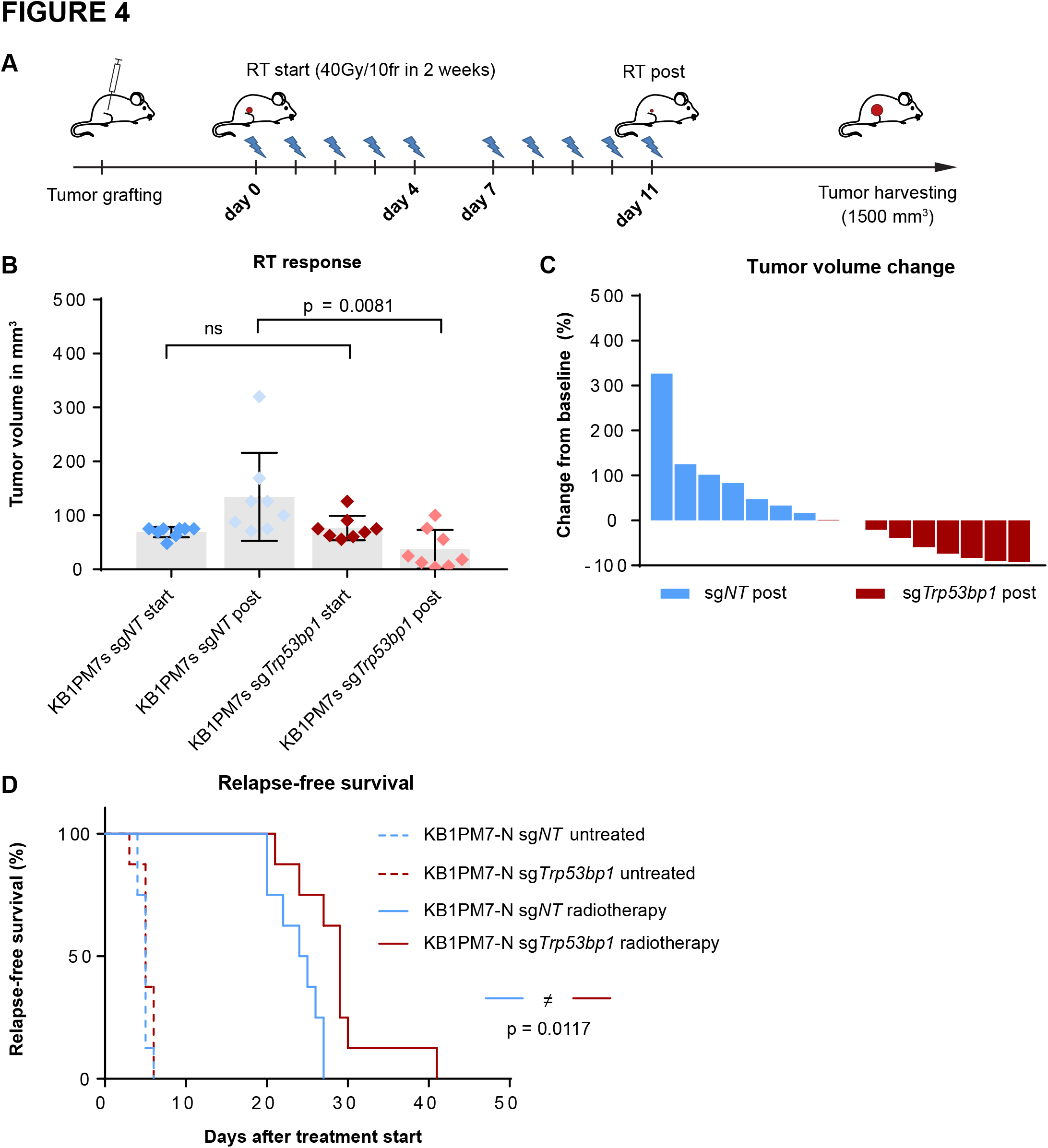
KB1PM7-N sg*Trp53bp1* targeted tumors show an enhanced radiotherapy response. **A.** Schematic overview depicting the RT treatment schedule. **B.** KB1PM7-N sg*NT* and KB1PM7-N sg*Trp53bp1* targeted tumor pieces were orthotopically transplanted in mice and were treated with radiotherapy (40Gy/10fr in 2 weeks) when tumors reached 50-100mm^3^. The tumor volume at the end of treatment was compared to the volume at the start of treatment. Data are plotted as mean ± SD. Significance was calculated by unpaired two-tailed students t-test. **C.** Waterfall plot showing the same data as in A, but plotted as relative change in tumor volume after treatment compared to treatment start. **D.** Kaplan-Meier curve showing that sg*Trp53bp1* targeted KB1PM7-N tumors have a prolonged time to relapse upon radiotherapy treatment, defined as five times the tumor volume compared to treatment initiation. Significance was calculated by Log-Rank (Mantel-Cox) test in Graphpad Prism.

## DISCUSSION

Acquired resistance to anticancer therapy may introduce new vulnerabilities, a concept known as collateral sensitivity (23). In this study, we demonstrate that this concept also applies to anticancer therapy targeting DDR-defects. We identified radiosensitivity as a therapeutically exploitable vulnerability of PARPi-resistant BRCA1-deficient cells that have restored HR via loss of the 53BP1 pathway. 53BP1 and its pathway member RIF1 have a known role in promoting DSB repair via NHEJ. However, while loss of 53BP1 or RIF1 in BRCA1-proficient cells has been associated with increased RT sensitivity (6,24), the interaction between 53BP1 and BRCA1 on RT response has remained unexplored. Collectively, our findings establish this interaction as a two-edged sword: loss of the 53BP1 pathway in BRCA1-deficient cells drives PARPi and topoisomerase inhibitor resistance at the expense of an acquired vulnerability to radiotherapy. These opposing responses to different DNA damaging agents might be reconciled by the context in which DNA damage arises: olaparib and topoisomerase inhibitors cause DSBs primarily during replication by inhibiting single-strand break repair and/or promoting PARP1/TOP1 trapping on the DNA, which ultimately results in replication fork collapse and the formation of one-sided DSBs (4). Therefore, a template for HR is in close proximity when a DSB arises, possibly explaining the strong effect of 53BP1 pathway inactivation on driving PARP- or topoisomerase inhibitor resistance. In contrast, acute DSB induction by RT is inflicted independent of the cell cycle and therefore more dependent on repair via the NHEJ pathway. Indeed, inhibition of NHEJ repair, for instance by knocking out KU70/80, was previously shown to sensitize cells to RT (25). We provide evidence that loss of the 53BP1 pathway also confers sensitivity to RT in a BRCA1-deficient context. Our data show that this acquired vulnerability can be exploited both *in vitro* and *in vivo* to constrain BRCA1-deficient cells that have acquired PARPi resistance via disruption of the 53BP1 pathway. Given the plethora of factors involved in the 53BP1 pathway and the pressing problem of clinical resistance to PARPi treatment, it will be important to determine the frequency of 53BP1/RIF1/REV7/Shieldin/CST pathway inactivation in PARPi-resistant tumors.

Our findings also re-emphasize the importance to understand the mechanistic basis of BRCA1-deficient tumors that have restored HR activity. For example, assessment of HR status by scoring for RAD51 IRIF formation may not provide sufficient detail to make informed treatment decisions since this assay does not discriminate BRCA1-dependent from BRCA1-independent restoration of HR. Given their hypersensitivity to radiotherapy, the latter tumors might have different treatment options from tumors in which BRCA1 expression has been restored.

## AUTHORS’ CONTRIBUTIONS

**Conception and design:** M. Barazas, G. Borst, J. Jonkers, S. Rottenberg

**Acquisition of data:** M. Barazas, A. Gasparini, Y. Huang, A. Kucukosmanoglu, S. Annunziato, W. Sol, A. Kersbergen, N. Proost, G. Borst, S. Rottenberg

**Analysis and interpretation of data:** M. Barazas, J. Jonkers, G. Borst, S. Rottenberg.

**Writing, review and/or revision of the manuscript:** M. Barazas, J. Jonkers, G. Borst, S. Rottenberg

**Technical, or material support:** P. Bouwman.

**Study supervision:** M. van de Ven, J. Jonkers, G. Borst, S. Rottenberg.

## ACKNOWLEDGMENTS

We wish to thank Piet Borst for critical reading of the manuscript, the members of the Preclinical Intervention Unit of the Mouse Clinic for Cancer and Aging (MCCA) at the Netherlands Cancer Institute (NKI) for their technical support with the animal experiments. We are grateful to the NKI animal facility, animal pathology facility and flow cytometry facility for their excellent service. Financial support came from the Dutch Cancer Society (KWF 2011-5220 and 2014-6532 to S.R. and J.J.), the Netherlands Organization for Scientific Research (VICI 91814643, NGI 93512009, Cancer Genomics Netherlands, and a National Roadmap Grant for Large-Scale Research Facilities to J.J.), the Swiss National Science Foundation (310030_156869 to S.R.), the Swiss Cancer League (KLS-4282-08-2017 to S.R.), ERC CoG-681572 to S.R., and ERC Synergy Grant 319661 to J.J.).

## REFERENCES

1. Nickoloff JA, Jones D, Lee SH, Williamson EA, Hromas R. Drugging the Cancers Addicted to DNA Repair. Journal of the National Cancer Institute 2017;109(11) doi 10.1093/jnci/djx059.

2. Knijnenburg TA, Wang L, Zimmermann MT, Chambwe N, Gao GF, Cherniack AD, et al. Genomic and Molecular Landscape of DNA Damage Repair Deficiency across The Cancer Genome Atlas. Cell Reports 2018;23(1):239–54.e6 doi 10.1016/j.celrep.2018.03.076.

3. Kaelin WG, Jr. The concept of synthetic lethality in the context of anticancer therapy. Nature reviews Cancer 2005;5(9):689–98 doi 10.1038/nrc1691.

4. Lord CJ, Ashworth A. PARP inhibitors: Synthetic lethality in the clinic. Science (New York, NY) 2017;355(6330):1152–8 doi 10.1126/science.aam7344.

5. Bunting SF, Callen E, Wong N, Chen HT, Polato F, Gunn A, et al. 53BP1 inhibits homologous recombination in Brca1-deficient cells by blocking resection of DNA breaks. Cell 2010;141(2):243–54 doi 10.1016/j.cell.2010.03.012.

6. Chapman JR, Barral P, Vannier JB, Borel V, Steger M, Tomas-Loba A, et al. RIF1 is essential for 53BP1-dependent nonhomologous end joining and suppression of DNA double-strand break resection. Mol Cell 2013;49(5):858–71 doi 10.1016/j.molcel.2013.01.002.

7. Jaspers JE, Kersbergen A, Boon U, Sol W, van Deemter L, Zander SA, et al. Loss of 53BP1 causes PARP inhibitor resistance in Brca1-mutated mouse mammary tumors. Cancer discovery 2013;3(1):68–81 doi 10.1158/2159-8290.CD-12-0049.

8. Xu G, Chapman JR, Brandsma I, Yuan J, Mistrik M, Bouwman P, et al. REV7 counteracts DNA double-strand break resection and affects PARP inhibition. Nature 2015;521(7553):541–4 doi 10.1038/nature14328.

9. Gupta R, Somyajit K, Narita T, Maskey E, Stanlie A, Kremer M, et al. DNA Repair Network Analysis Reveals Shieldin as a Key Regulator of NHEJ and PARP Inhibitor Sensitivity. Cell 2018;173(4):972–88.e23 doi 10.1016/j.cell.2018.03.050.

10. Barazas M, Annunziato S, Pettitt SJ, de Krijger I, Ghezraoui H, Roobol SJ, et al. The CST Complex Mediates End Protection at Double-Strand Breaks and Promotes PARP Inhibitor Sensitivity in BRCA1-Deficient Cells. Cell Rep 2018;23(7):2107–18 doi 10.1016/j.celrep.2018.04.046.

11. Liu X, Holstege H, van der Gulden H, Treur-Mulder M, Zevenhoven J, Velds A, et al. Somatic loss of BRCA1 and p53 in mice induces mammary tumors with features of human BRCA1-mutated basal-like breast cancer. Proceedings of the National Academy of Sciences of the United States of America 2007;104(29):12111–6 doi 10.1073/pnas.0702969104.

12. Rottenberg S, Jaspers JE, Kersbergen A, van der Burg E, Nygren AO, Zander SA, et al. High sensitivity of BRCA1-deficient mammary tumors to the PARP inhibitor AZD2281 alone and in combination with platinum drugs. Proceedings of the National Academy of Sciences of the United States of America 2008;105(44):17079–84 doi 10.1073/pnas.0806092105.

13. Zander SA, Kersbergen A, van der Burg E, de Water N, van Tellingen O, Gunnarsdottir S, et al. Sensitivity and acquired resistance of BRCA1;p53-deficient mouse mammary tumors to the topoisomerase I inhibitor topotecan. Cancer research 2010;70(4):1700–10 doi 10.1158/0008-5472.Can-09-3367.

14. Duarte AA, Gogola E, Sachs N, Barazas M, Annunziato S, J RdR, et al. BRCA-deficient mouse mamm ary tumor organoids to study cancer-drug resistance. Nat Methods 2017 doi 10.1038/nmeth.4535.

15. Cong L, Ran FA, Cox D, Lin S, Barretto R, Habib N, et al. Multiplex genome engineering using CRISPR/Cas systems. Science (New York, NY) 2013;339(6121):819–23 doi 10.1126/science.1231143.

16. Harmsen T, Klaasen S, van de Vrugt H, Te Riele H. DNA mismatch repair and oligonucleotide end-protection promote base-pair substitution distal from a CRISPR/Cas9-induced DNA break. Nucleic Acids Res 2018;46(6):2945–55 doi 10.1093/nar/gky076.

17. Chen J, Silver DP, Walpita D, Cantor SB, Gazdar AF, Tomlinson G, et al. Stable interaction between the products of the BRCA1 and BRCA2 tumor suppressor genes in mitotic and meiotic cells. Mol Cell 1998;2(3):317–28.

18. Follenzi A, Ailles LE, Bakovic S, Geuna M, Naldini L. Gene transfer by lentiviral vectors is limited by nuclear translocation and rescued by HIV-1 pol sequences. Nat Genet 2000;25(2):217–22 doi 10.1038/76095.

19. Sakaue-Sawano A, Kurokawa H, Morimura T, Hanyu A, Hama H, Osawa H, et al. Visualizing spatiotemporal dynamics of multicellular cell-cycle progression. Cell 2008;132(3):487–98 doi 10.1016/j.cell.2007.12.033.

20. Prahallad A, Heynen GJ, Germano G, Willems SM, Evers B, Vecchione L, et al. PTPN11 Is a Central Node in Intrinsic and Acquired Resistance to Targeted Cancer Drugs. Cell Rep 2015;12(12):1978–85 doi 10.1016/j.celrep.2015.08.037.

21. Brinkman EK, Chen T, Amendola M, van Steensel B. Easy quantitative assessment of genome editing by sequence trace decomposition. Nucleic Acids Res 2014;42(22):e168 doi 10.1093/nar/gku936.

22. Clarkson R, Lindsay PE, Ansell S, Wilson G, Jelveh S, Hill RP, et al. Characterization of image quality and image-guidance performance of a preclinical microirradiator. Medical physics 2011;38(2):845–56 doi 10.1118/1.3533947.

23. Hutchison DJ. CROSS RESISTANCE AND COLLATERAL SENSITIVITY STUDIES IN CANCER CHEMOTHERAPY. Advances in cancer research 1963;7:235–50.

24. Nakamura K, Sakai W, Kawamoto T, Bree RT, Lowndes NF, Takeda S, et al. Genetic dissection of vertebrate 53BP1: a major role in non-homologous end joining of DNA double strand breaks. DNA repair 2006;5(6):741–9 doi 10.1016/j.dnarep.2006.03.008.

25. Smith GC, Jackson SP. The DNA-dependent protein kinase. Genes Dev 1999;13(8):916–34.

